# Inhibition of Soluble Epoxide Hydrolase Reduces Inflammation and Myocardial Injury in Arrhythmogenic Cardiomyopathy

**DOI:** 10.1101/2024.02.17.580812

**Authors:** Dipak Panigrahy, Abigail G. Kelly, Weicang Wang, Jun Yang, Sung Hee Hwang, Michael Gillespie, Isabella Howard, Carlos Bueno-Beti, Angeliki Asimaki, Vinay Penna, Kory Lavine, Matthew L. Edin, Darryl C. Zeldin, Bruce D. Hammock, Jeffrey E. Saffitz

## Abstract

Previous studies have implicated persistent innate immune signaling in the pathogenesis of arrhythmogenic cardiomyopathy (ACM), a familial non-ischemic heart muscle disease characterized by life-threatening arrhythmias and progressive myocardial injury. Here, we provide new evidence implicating inflammatory lipid autocoids in ACM. We show that specialized pro-resolving lipid mediators are reduced in hearts of *Dsg2^mut/mut^* mice, a well characterized mouse model of ACM. We also found that ACM disease features can be reversed in rat ventricular myocytes expressing mutant *JUP* by the pro-resolving epoxy fatty acid (EpFA) 14,15-eicosatrienoic acid (14-15-EET), whereas 14,15-EE-5(Z)E which antagonizes actions of the putative 14,15-EET receptor, intensified nuclear accumulation of the desmosomal protein plakoglobin. Soluble epoxide hydrolase (sEH), an enzyme that rapidly converts pro-resolving EpFAs into polar, far less active or even pro-inflammatory diols, is highly expressed in cardiac myocytes in *Dsg2^mut/mut^* mice. Inhibition of sEH prevented progression of myocardial injury in *Dsg2^mut/mut^* mice and led to recovery of contractile function. This was associated with reduced myocardial expression of genes involved in the innate immune response and fewer pro-inflammatory macrophages expressing CCR2, which mediate myocardial injury in *Dsg2^mut/mut^* mice. These results suggest that pro-inflammatory eicosanoids contribute to the pathogenesis of ACM and, further, that inhibition of sEH may be an effective, mechanism-based therapy for ACM patients.

## INTRODUCTION

Arrhythmogenic cardiomyopathy (ACM) is a familial non-ischemic heart muscle disease characterized by arrhythmias and progressive myocardial injury, typically involving the right ventricle.^1,2^ Most cases are caused by pathogenic variants in genes that encode desmosomal proteins.^1,2^ We have previously reported that inhibition of nuclear factor κB (NFκB) signaling rescues the disease phenotype and reduces myocardial expression of pro-inflammatory cytokines in a well-characterized mouse model of ACM involving homozygous knock-in of a variant in the gene for the desmosomal protein, desmoglein-2 (*Dsg2^mut/mut^* mice).^3,4^ NFκB signaling is also activated *in vitro* in induced pluripotent stem cell (iPSC)-cardiac myocytes derived from ACM patients with disease-causing variants in *PKP*2 or *DSG2,*^3,5^ and in rat ventricular myocytes expressing a disease-related variant in *JUP* (all genes that encode desmosomal proteins).^3^ ACM iPSC-cardiac myocytes and rat myocytes expressing mutant *JUP* produce and secrete large amounts of pro-inflammatory mediators under basal conditions without prior stimulation or provocation.^3,5–7^ Taken together, these observations suggest that the pathogenesis of ACM is driven by a persistent, cardiac myocyte-autonomous innate immune response that fails to resolve.

Here, we show that levels of specialized pro-resolving mediators (SPMs)^8^ are reduced in hearts of *Dsg2^mut/mut^* mice compared to wild type (WT) mice. Hearts of *Dsg2^mut/mut^* mice also show increased expression of genes activated in endoplasmic reticulum stress (ER stress), a hallmark of unresolved inflammation.^9^ In addition, ACM disease features are reversed in rat myocytes expressing mutant *JUP* by the pro-resolving epoxy fatty acid (EpFA) 14,15-EET, whereas 14,15-EE-5(Z)E (a structural analog of 14,15-EET), which antagonizes actions of the putative 14,15-EET receptor, intensifies nuclear accumulation of the desmosomal protein plakoglobin (aka, γ-catenin), which has been implicated in disease pathogenesis in ACM patients.^10^ Soluble epoxide hydrolase (sEH), an enzyme responsible for metabolizing pro-resolving EpFAs into pro-inflammatory diols,^11,12^ is highly expressed in cardiac myocytes in hearts of WT and *Dsg2^mut/mut^* mice, confirming previous studies.^13^ Inhibition of sEH prevents progression of the disease in *Dsg2^mut/mut^* mice and promotes significant recovery of contractile function associated with reduced myocardial injury, diminished expression of genes activated in the innate immune response, and fewer pro-inflammatory macrophages expressing CCR2. sEH inhibitors are currently being evaluated in early stage clinical safety trials in patients with chronic neuropathic pain as a clinical path^14^ Our results suggest that inhibition of sEH may be a novel mechanism-based therapy for ACM patients.

## METHODS

### *In vivo* and *in vitro* models of ACM

All animal studies were in full compliance with policies of Beth Israel Deaconess Medical Center and St. George’s University of London, and conformed to the *Guide for the Care and Use of Laboratory Animals* from the National Institutes of Health (NIH publication no. 85-23, revised 1996). Mice for *in vivo* studies were housed in a 12-hour-light/dark cycle, climate-controlled facility with *ad libitum* access to water and standard rodent chow. Studies were performed in wild type (WT) mice and mice with homozygous knock-in of a variant in *Dsg2*, the gene encoding the desmosomal cadherin desmoglein-2 (*Dsg2^mut/mut^* mice) as previously described.^3,4,6,7^ This variant entails loss of exons 4 and 5 which causes a frameshift and premature termination of translation.

*In vitro* studies were performed in primary cultures of ventricular myocytes prepared from disaggregated ventricles of 1-day-old Wistar rat pups as previously described.^3,6,7^ Cells were plated on collagen-coated plastic chamber slides at a density of 2.4 x 10^5^ cells/cm^2^. Two days post-plating, monolayers were transfected in serum-free medium for 1hour with a recombinant adenoviral construct (pAd/CMV/V5-DEST vector) containing 2157del2 *JUP*, after which the viral solution was replaced with complete medium. 24 hours later, cultures were incubated with 10 μM 14,15-epoxyeicosatrieonic acid (14,15-EET) or 10 μM 14,15- epoxyeicosa-5(Z)-enoic acid (14,15-EEZE) for an additional 24 hours. Other cultures were incubated for 24 hours with 1 µM TPPU (1-trifluoro-methoxy-phenyl-3-(1-propionylpiperidin-4- yl) urea), a small molecule inhibitor of sEH,^16^ or 500 nM PTUPB (4-(5-phenyl-3-{3-[3-(4- trifluoromethylphenyl)-ureido]-propyl}-pyrazol-1-yl)-benzenesulfonamide), a dual eicosanoid pathway (COX2/sEH) inhibitor.^16^ Transfected cultures treated with vehicle only and non-transfected cultures were used as controls. Thereafter, the cultures were rinsed in serum-free medium and fixed in 4% paraformaldehyde at 25°C for 5 minutes in preparation for immunofluorescence microscopy as previously described.^3,6,7^ Fixed cells were incubated with mouse monoclonal anti-plakoglobin (Sigma), anti-Cx43 (Millipore) and anti-RelA/p65 (LSBio) antibodies. Secondary antibodies included Cy3-conjugated goat anti-mouse or anti-rabbit IgGs (H+L; Jackson Immunoresearch). The cells were counterstained with DAPI and examined by laser scanning confocal microscopy (Nikon A1).

### Inhibition of sEH and characterization of disease phenotypes in *Dsg2^mut/mut^* mice

The effects of the small molecule sEH inhibitor TPPU were studied in age-matched WT and *Dsg2^mut/mut^* mice. 9-week-old animals underwent echocardiography before being implanted with intraperitoneal osmotic mini-pumps (Alzet, Model 1004) as previously described.^3,4,16^ They received either vehicle or drug (TPPU dissolved in DMSO/polyethylene glycol). Drug-treated mice received 5mg/kg/day TPPU via continuous infusion (1µL/hour for 28 days); vehicle-treated mice received an equivalent volume of vehicle for 28 days. Final echocardiograms were obtained in both groups at 13 weeks of age. Thereafter, animals were euthanized and hearts were collected for additional studies including histology to measure the amount of ventricular fibrosis; immunohistochemistry to measure the number of CCR2+ cells; western blotting and qPCR to measure amounts of specific proteins and gene expression levels; and ELISA to quantify specific lipid mediators in hearts. Western blots were performed on lysates of hearts from WT and *Dsg2^mut/mut^* mice using methods described in previous studies.^17,18^ ELISA assays were performed on extracts of hearts from WT and *Dsg2^mut/mut^* mice using kits and protocols from Cayman Chemicals as described in previous studies.^17,19^ qPCR was used to measure expression of genes related to the ER-stress response and innate immune signaling as described in previous studies.^18^

### Statistical analysis

All data are presented as mean ± SEM; n-values and the statistical analyses performed for each experiment are indicated in figure legends. Differences in measured variables were assessed with Mann-Whitney or multiple comparisons ANOVA with Tukey post-hoc analysis. A p-value of <0.05 was considered statistically significant. All statistical analyses were analyzed using GraphPad Prism (v9.2) software.

## RESULTS

### *Dsg2^mut/mut^* mice have reduced levels of pro-resolving lipid mediators and express markers of endoplasmic reticulum (ER) stress

We used ELISA to quantify levels of selected pro-resolving lipid mediators in hearts of *Dsg2^mut/mut^* mice. Levels of the specialized pro-resolving mediators (SPMs) resolvin D1, resolvin D2 and maresin-1 were all significantly reduced in hearts of *Dsg2^mut/mut^* mice compared to WT mice, as was expression of GRP18, the receptor for resolvin D2 measured by western blotting (**Figure 1A**). ER stress is a hallmark of unresolved innate immune signaling.^9^ As shown in **Figure 1B**, expression of the ER stress response genes *BiP*, which acts as an ER chaperone, and *Pdi,* involved in protein folding, was increased in hearts of *Dsg2^mut/mut^* mice compared to WT mice. Taken together, these results provide evidence of persistent inflammation in *Dsg2^mut/mut^* mice that fails to resolve.

**Figure 1:**
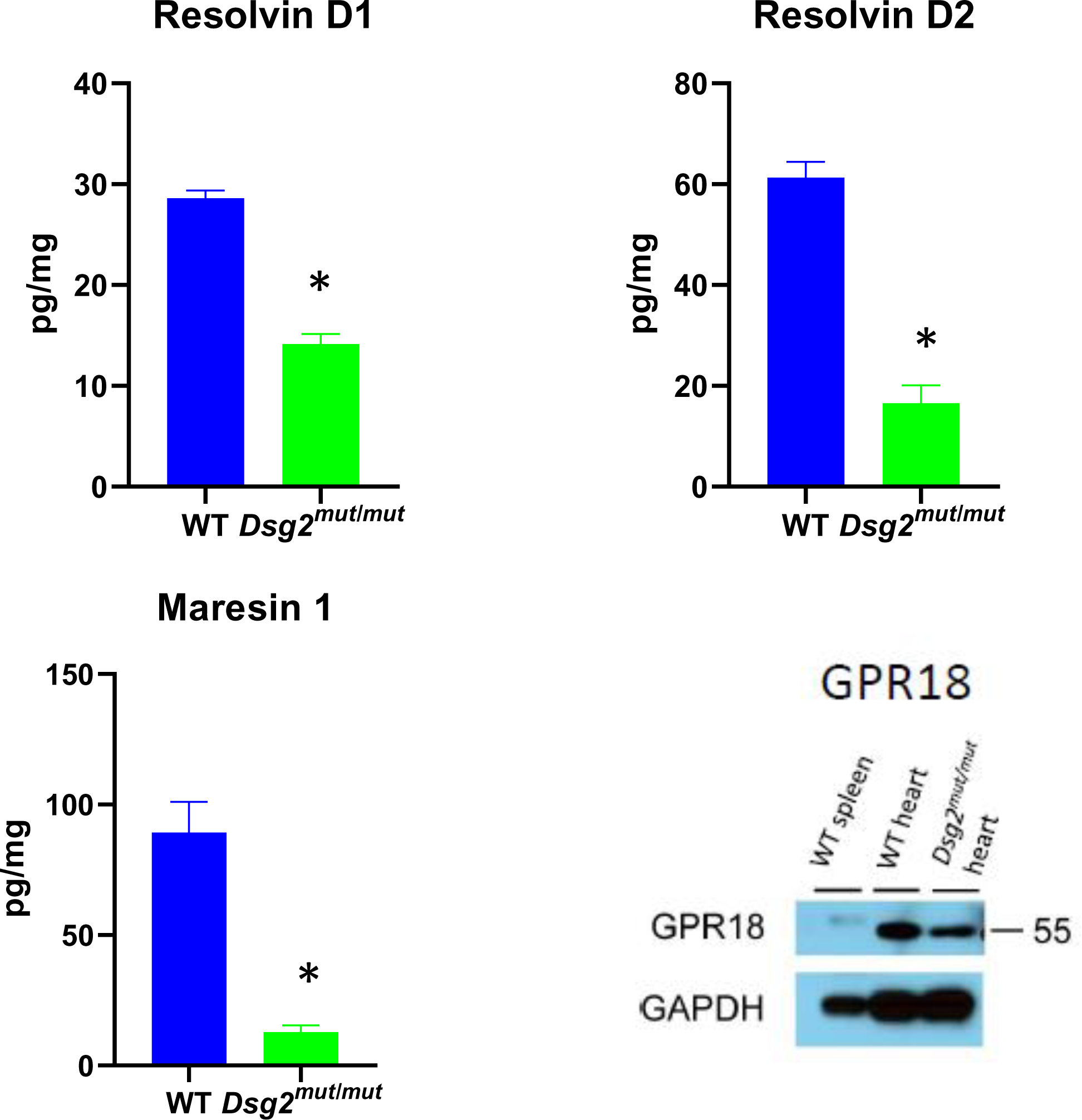
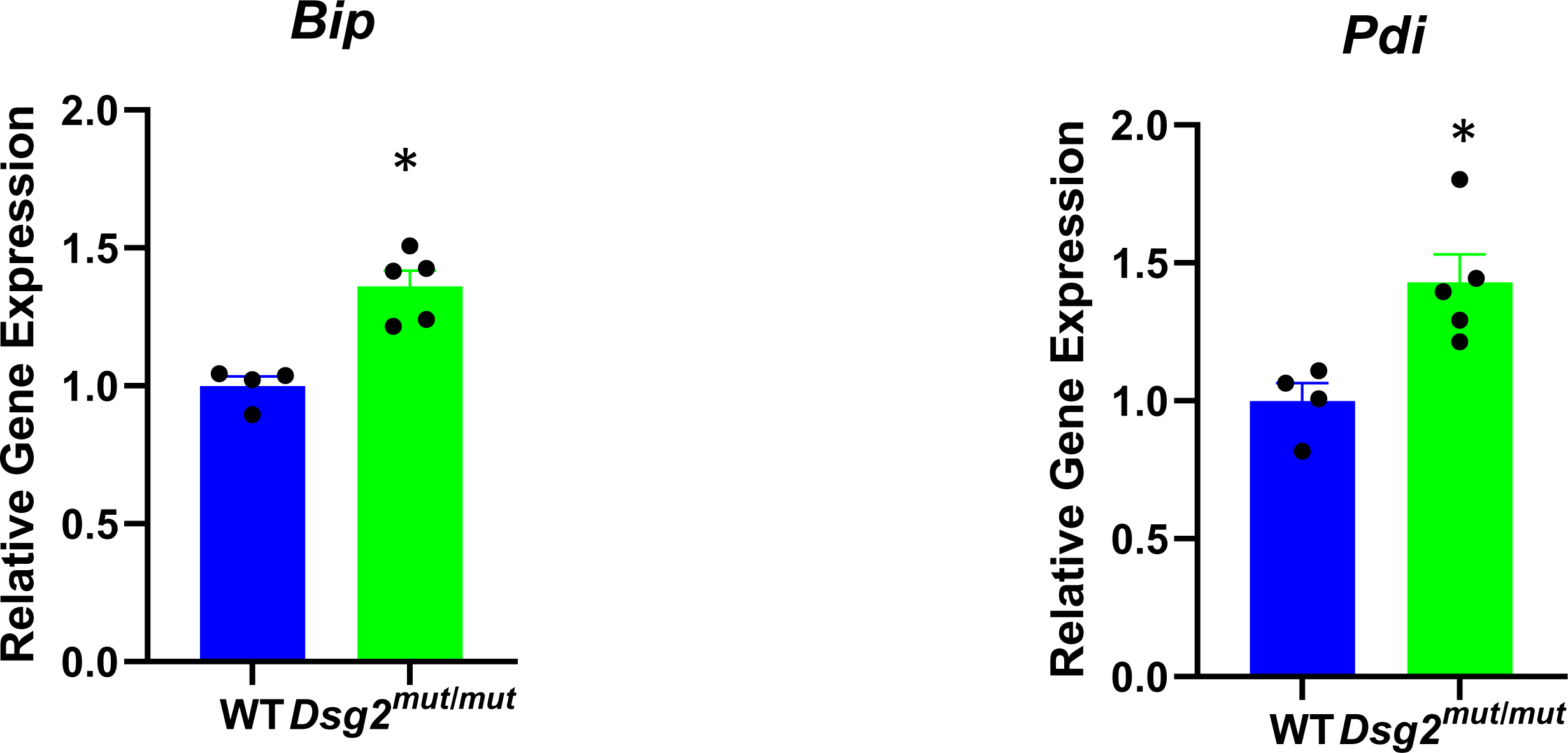
**A.** Reduced levels of specialized pro-resolving mediators resolvin D1, maresin 1 and resolvin E2 in hearts of 16-week-old *Dsg2^mut/mut^* compared to wildtype (WT) mice measured by ELISA, and of GPR18 in hearts of 16-week-old *Dsg2^mut/mut^*measured by western blotting. A sample of spleen was used as a positive control in the western blot; * p<0.05 vs. WT by Mann Whitney U test. **B.** qPCR showing increased expression of the endoplasmic reticulum chaperone gene *BiP* and the protein folding protein disulfide isomerase gene *Pdi*, both markers of endoplasmic reticulum stress. Gene expression values in wildtype samples were normalized to 1; values in *Dsg2^mut/mut^* samples are shown as relative levels; * p<0.05 vs. WT by Mann Whitney U test.

### 14,15-EET blocks NFκB signaling and rescues ACM disease features in vitro

We have previously reported that NFκB signaling is activated in a well-characterized *in vitro* model of ACM involving neonatal rat ventricular myocytes that express a variant in *JUP*, the gene for the desmosomal protein plakoglobin.^3^ These cells exhibit characteristic features seen in ACM patients including redistribution of junctional plakoglobin to intracellular and nuclear sites and loss of cell surface signal for Cx43, the major ventricular gap junction protein.^3,6,7^ They also exhibit activation of NFκB signaling under basal conditions *in vitro*.^3^ As shown in **Figure 2**, junctional plakoglobin and Cx43 distribution was normalized when cells were incubated with the pro-resolving EpFA 14,15-eicosatrienoic acid (14,15-EET). In addition, NFκB signaling, indicated by the presence of nuclear signal for phospho-RelA/p65, was turned off in cells exposed to 14,15-EET. By contrast, nuclear accumulation of plakoglobin (aka γ-catenin) was greatly increased in cells incubated with 10µM 14,15-EE-5(Z)E, which antagonizes actions of 14,15-EET (**Figure 2**). This was of particular interest, as nuclear translocation of plakoglobin in cardiac myocytes is typically seen in patients with ACM,^20^ and has been implicated in altered Wnt signaling in the pathogenesis of ACM.^10^

**Figure 2.**
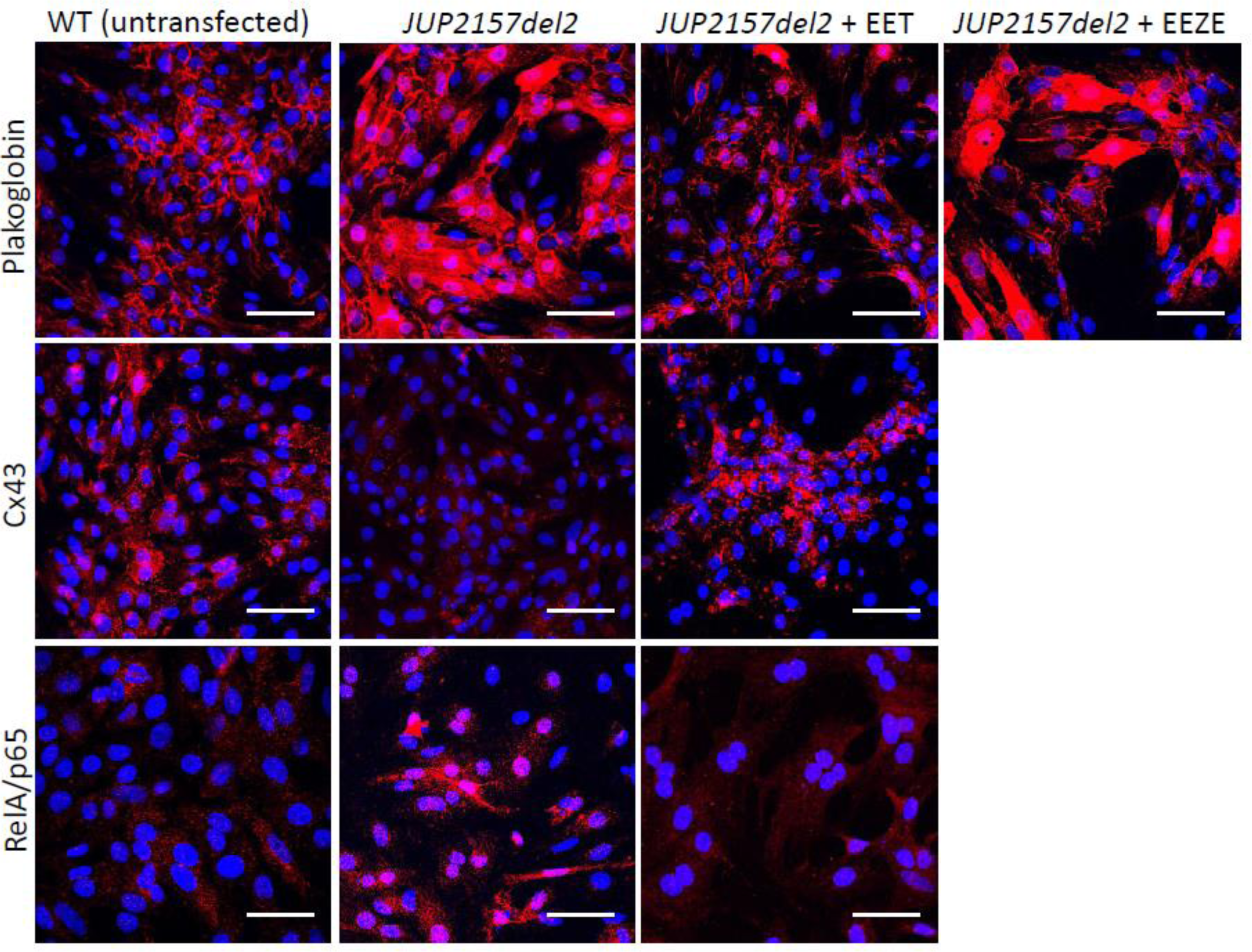
Effects of 14,15-eicosatrienoic acid (14,15-EET) and 14,15-eicosa-5(Z)-enoic acid (14,15-EEZE) on the distribution of immunoreactive signals for plakoglobin, connexin43 (Cx43) and RelA/p65 in primary cultures of wildtype (WT) neonatal rat ventricular myocytes and myocyes transfected to express the ACM disease allele *JUP2157del2*.

### sEH is expressed mainly in cardiac myocytes in hearts of *Dsg2^mut/mut^* mice

We previously performed single nucleus RNA sequencing (sn-RNAseq) in cells isolated from hearts of 16-week-old *Dsg2^mut/mut^* and WT mice.^4^ Here, we analyzed the sn-RNAseq data to identify the specific cell types that expressed *Ephx2*, the major sEH gene in mice. We found that in and *Dsg2^mut/mut^* mouse hearts, sEH is expressed mainly in cardiac myocytes and to a lesser extent in endothelial and myeloid cells (**Figure 3)**. Similar data were seen in WT mouse hearts. These observations are in accordance with and confirm a recent report showing that selective disruption of sEH in cardiac myocytes (but not endothelial cells) improves recovery following ischemia/reperfusion injury in mouse hearts.^13^

**Figure 3.**
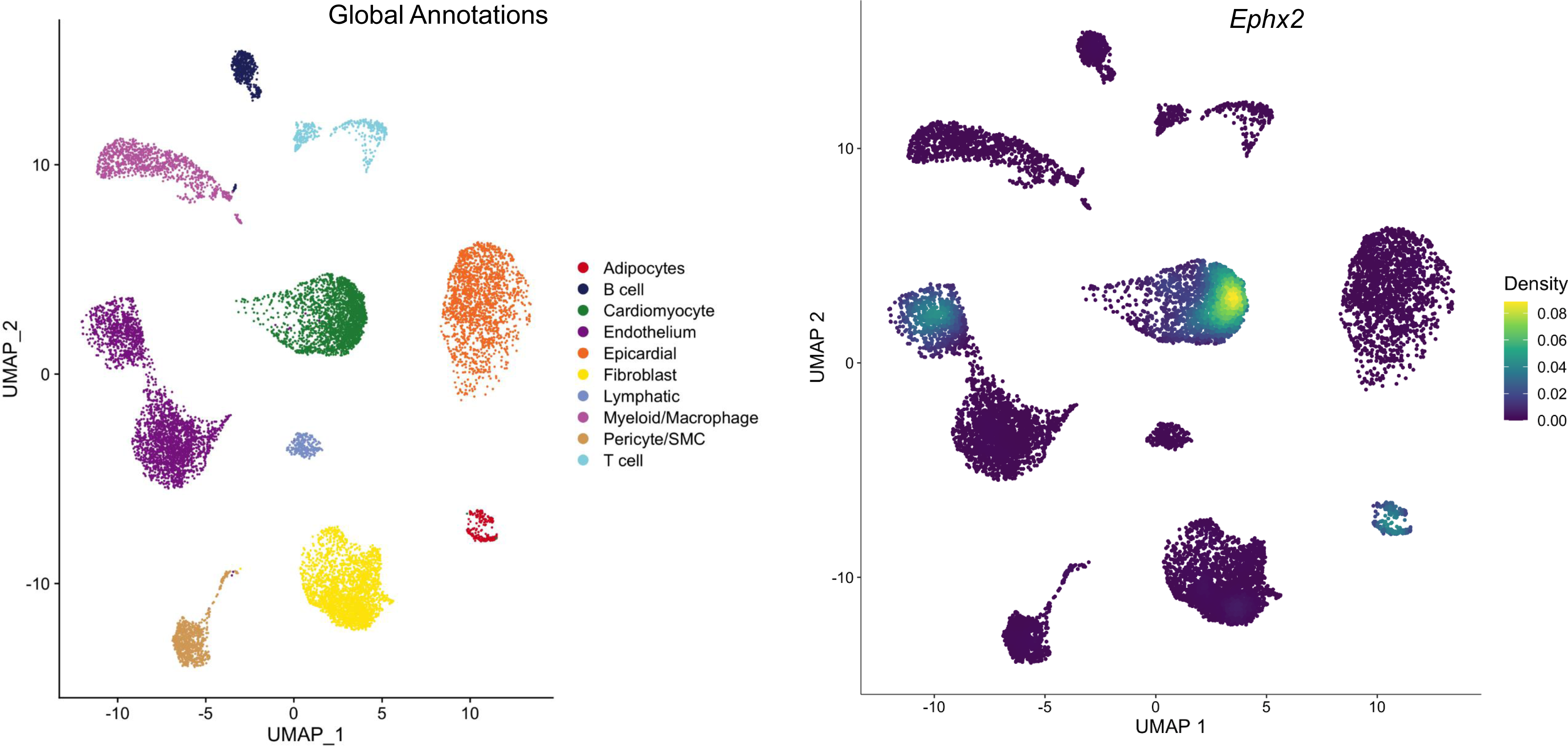
UMAPs showing populations of cells isolated from hearts of 16-week-old *Dsg2^mut/mut^* mice and the relative density of *Ephx2* expression in each population.

### Inhibition of sEH normalizes ACM disease features *in vitro* and prevents myocardial injury in *Dsg2^mut/mut^* mice

We have previously reported that inhibition of NFκB corrects ACM disease features in cultured rat myocytes expressing mutant *JUP*.^3^ It also rescues the disease phenotype in *Dsg2^mut/mut^* mice.^3,4^ To determine if inhibition of sEH also mitigates the ACM disease phenotype and turns off NFκB signaling *in vitro*, we incubated cultures of rat myocytes expressing *JUP2157del2* with the sEH inhibitor TPPU or the dual COX-2/sEH inhibitor PTUPB. As shown in **Figure 4**, TPPU and PTUPB both restored the normal cell surface distribution of plakoglobin and Cx43 in rat myocytes expressing mutant *JUP* and eliminated nuclear signal for RelA/p65 indicating that NFκB signaling was reduced. Thus, inhibition of sEH in primary cultures of cardiac myocytes *in vitro* reverses characteristic features of the ACM disease phenotype seen in patients. These data further suggest that cyclooxygenase inhibitors can synergize with sEH inhibitors in reversing disease features in ACM and support previous observations regarding multiple pro-inflammatory pathways in ACM.^3,4^

**Figure 4.**
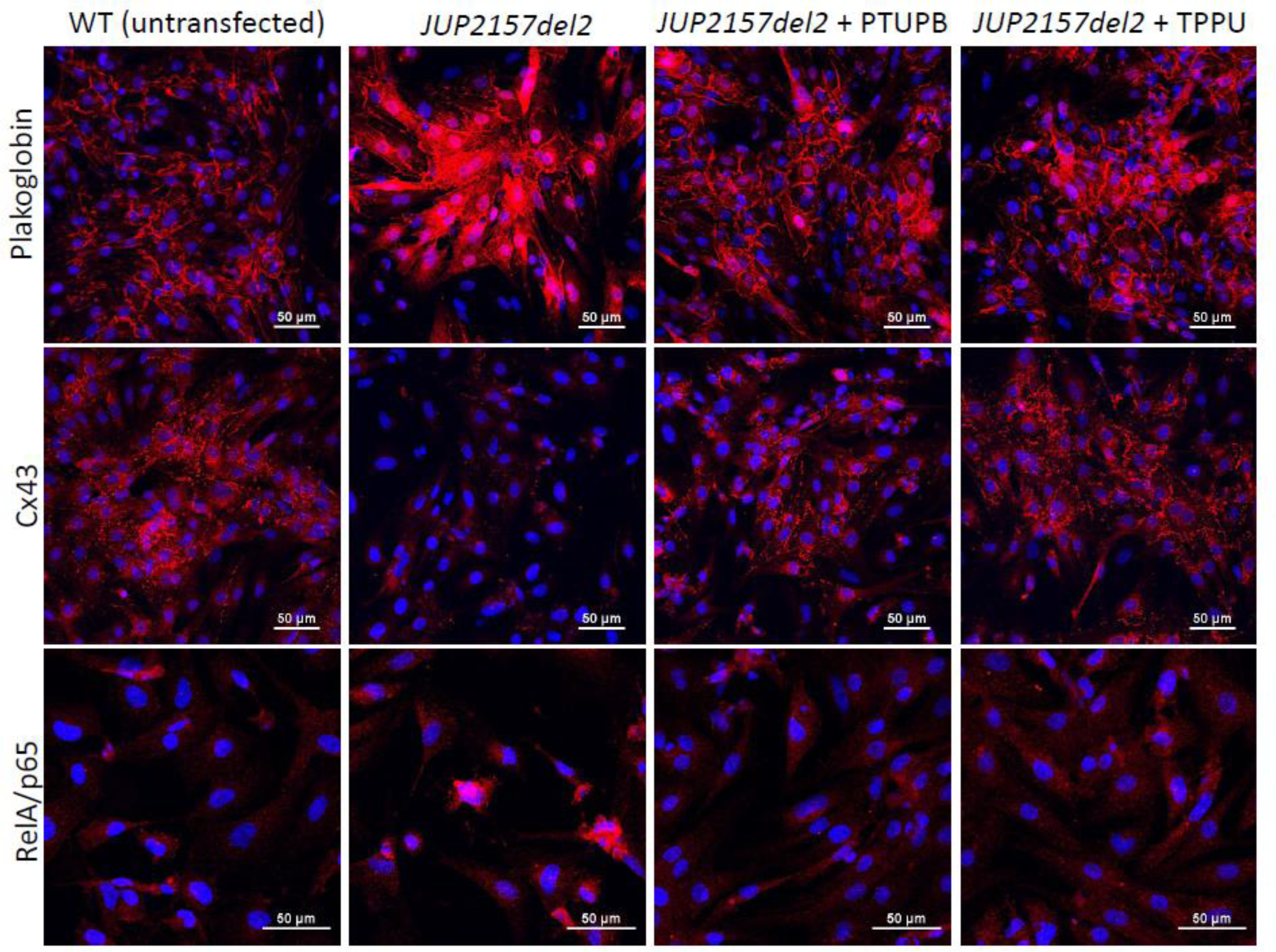
Effects of 1 µM 1-trifluoro-methoxy-phenyl-3-(1-propionylpiperidin-4-yl) urea (TPPU) and 500 nM 4-(5-phenyl-3-{3-[3-(4-trifluoromethylphenyl)-ureido]-propyl}-pyrazol-1- yl)-benzenesulfonamide (PTUPB) on the distribution of of immunoreactive signals for plakoglobin, connexin43 (Cx43) and RelA/p65 in primary cultures of wildtype (WT) neonatal rat ventricular myocytes and myocyes transfected to express the ACM disease allele *JUP2157del2*.

To determine if inhibition of sEH can promote resolution of inflammation and limit myocardial injury *in vivo*, we treated 9-week old *Dsg2^mut/mut^* mice for 4 weeks with TPPU and monitored effects on left ventricular contractile function and immune signaling. As shown in **Figure 5**, disease progressed during the 4-week treatment interval in *Dsg2^mut/mut^* mice given vehicle. These mice exhibited significant deterioration of LV ejection fraction and reduced LV fractional shortening. By contrast, *Dsg2^mut/mut^* mice treated with TPPU showed significant improvement in left ventricular ejection fraction and LV fractional shortening to levels roughly equivalent to those seen in age-matched WT mice. Every treated *Dsg2^mut/mut^* mouse showed enhanced contractile performance whereas untreated *Dsg2^mut/mut^* mice showed no improvement or exhibited functional deterioration (**Figure 5**). TPPU had no effect on left ventricular contractile function in WT mice. As previously reported, *Dsg2^mut/mut^* mice show little if any fibrosis at ∼8 weeks of age.^3,4,6,7^ However, marked ventricular fibrosis was seen in 13-week vehicle-treated *Dsg2^mut/mut^* mice, whereas significantly less fibrosis was present in hearts of *Dsg2^mut/mut^* mice treated with TPPU (**Figure 6A**). Similarly, as previously reported,^4^ the number of cells expressing CCR2 was increased by ∼5-fold in hearts of untreated *Dsg2^mut/mut^* mice, but they were significantly reduced in number in hearts of treated *Dsg2^mut/mut^* mice (**Figure 6B**).

**Figure 5.**
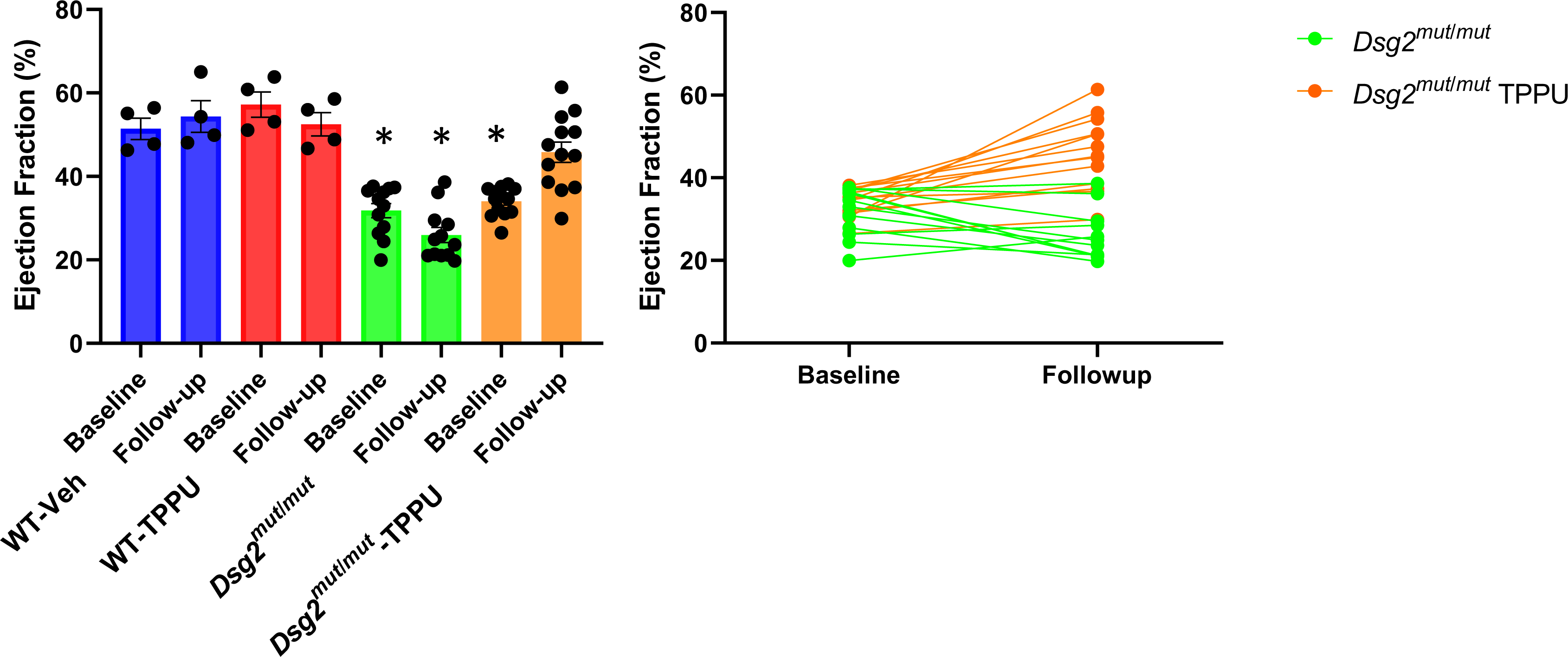
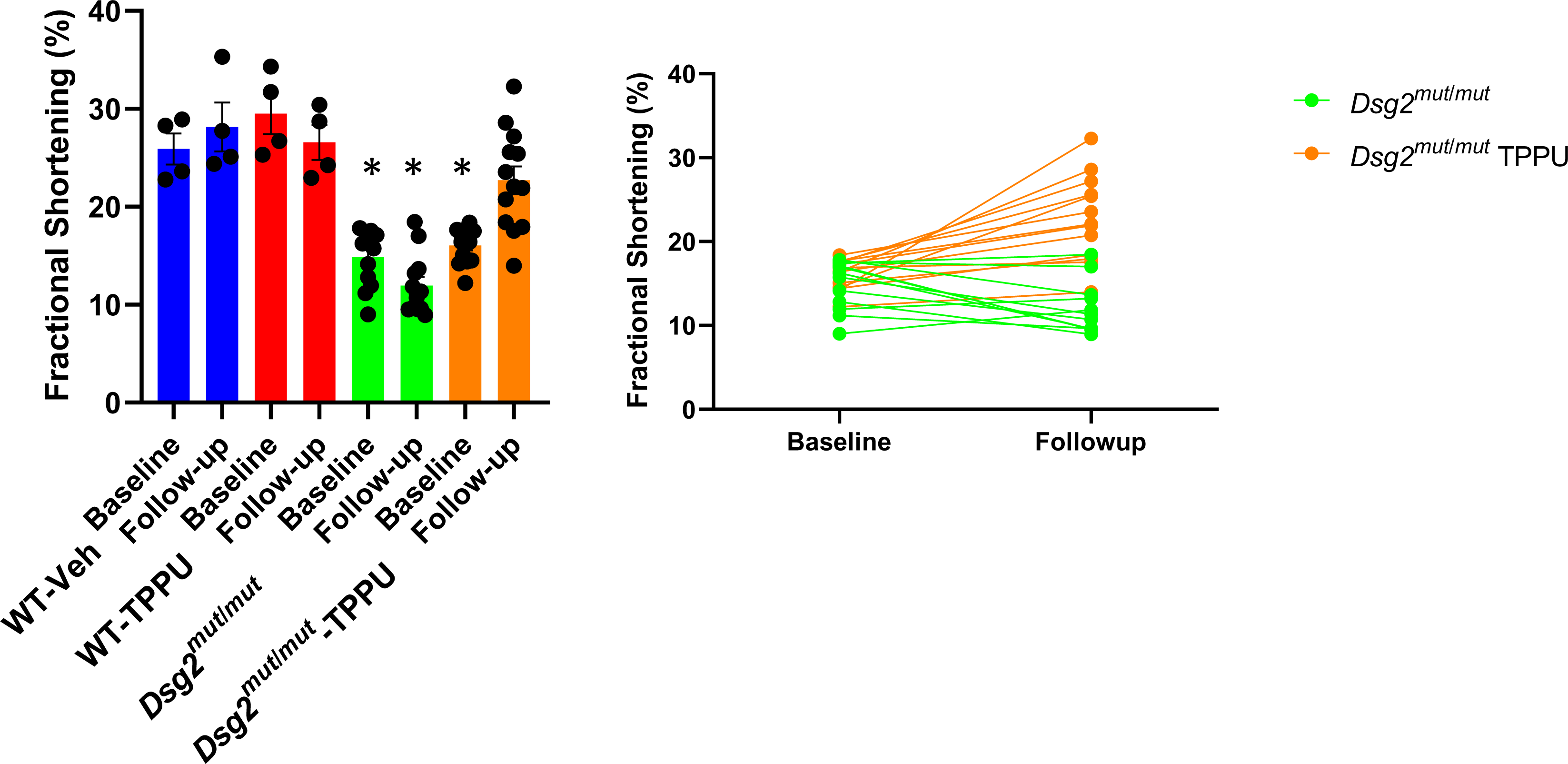
Effects of 1-trifluoro-methoxy-phenyl-3-(1-propionylpiperidin-4-yl) urea (TPPU) on left ventricular ejection fraction (**A**) and fractional shortening (**B**) in wildtype (WT) and *Dsg2^mut/mut^* mice. Baseline echocardiography was performed in 9-week-old mice and then repeated after treatment for 4 weeks with TPPU or vehicle (Veh). Data are shown for each group (left) and each individual animal (right); * p<0.0001 vs. WT-Veh by multiple comparisons ANOVA.

**Figure 6.**
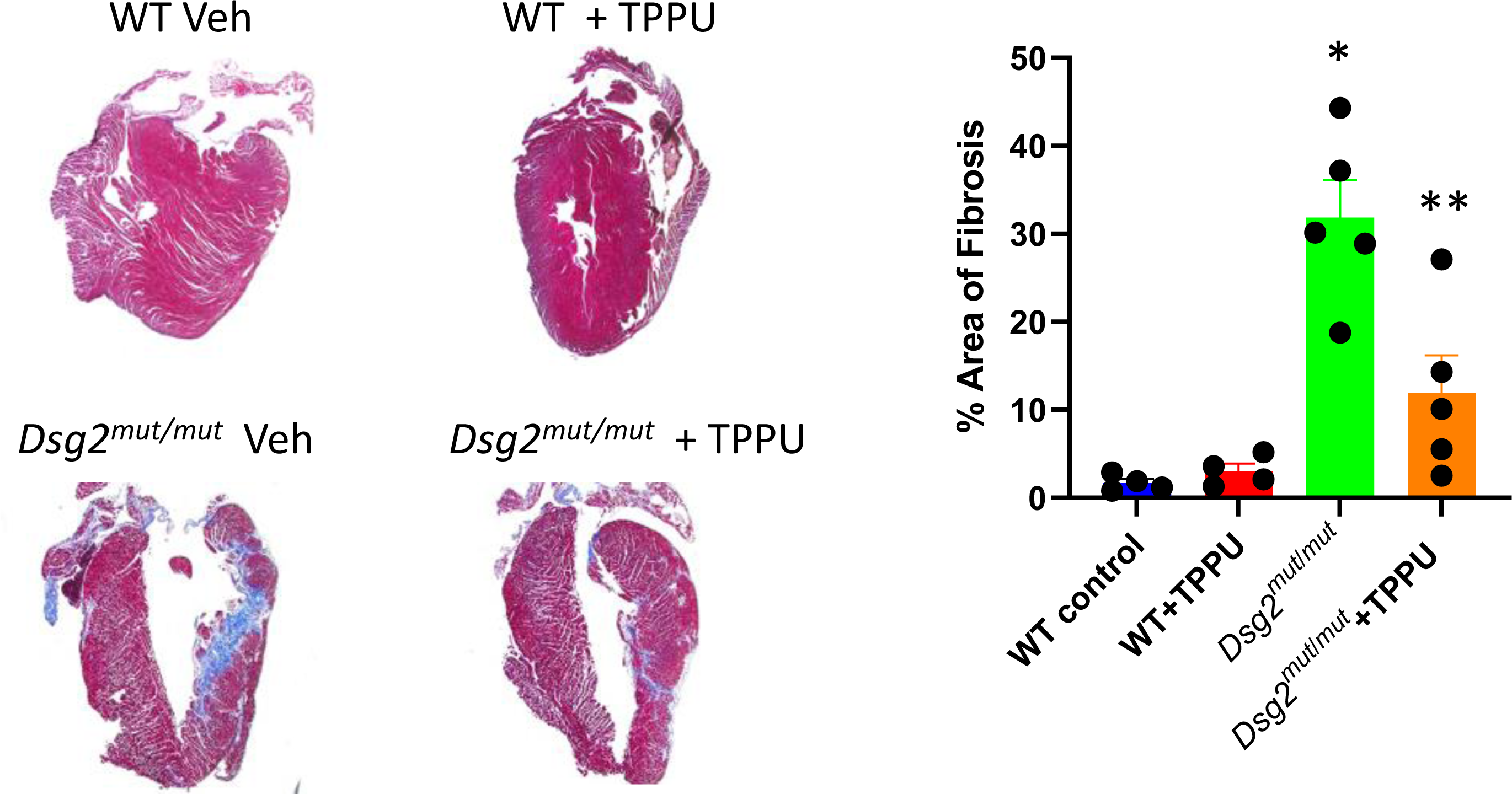
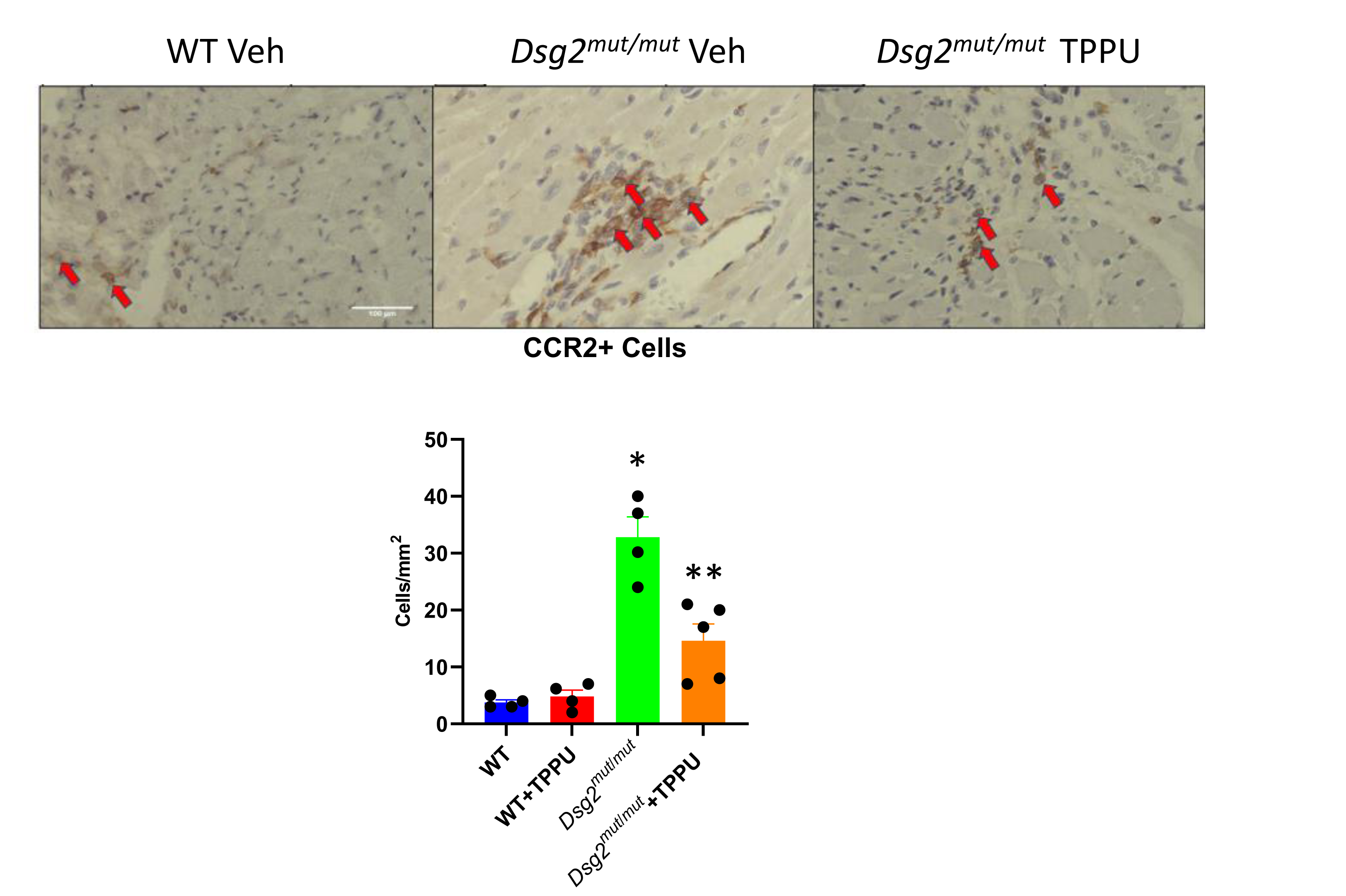
Effects of 1-trifluoro-methoxy-phenyl-3-(1-propionylpiperidin-4-yl) urea (TPPU) on the amount of myocardial fibrosis (**A**) and the number of cells expressing CCR2 (**B**) in wildtype (WT) and *Dsg2^mut/mut^* mice (positive cells identified by arrows in representative immunostained tissue sections). Hearts were excised from animals after treatment for 4 weeks with TPPU or vehicle (Veh) and analyzed by histology in trichrome stained sections or by immunohistochemistry in sections stained with an anti-CCR2 antibody; * p<0.05 vs. WT; ** p<0.05 vs. vehicle-treated *Dsg2^mut/mut^* mice by multiple T test.

These observations are of particular interest as we have shown that pro-inflammatory myeloid cells expressing CCR2 are highly injurious to the heart in ACM.^4^ Lastly, we have previously reported increased expression of key genes involved in the innate immune response in hearts of *Dsg2^mut/mut^* mice including *Tnfα* (the gene for the pro-inflammatory cytokine tumor necrosis factor-alpha), *Tlr4* (which encodes Toll-like receptor 4, the major pattern recognition receptor on cardiac myocytes), and *Ccr2* which is expressed by pro-inflammatory macrophages.^3,4^ These findings reflect a state of persistent innate immune signaling. As shown in **Figure 7**, we confirmed upregulation of these genes in *Dsg2^mut/mut^* mice and, further, showed that inhibiton of sEH reduced myocardial expression of these genes. Taken together with data in **Figure 6**, these results show that inhibition of sEH promotes resolution of inflammation leading to significant functional recovery and cessation of ACM disease progression.

**Figure 7.**
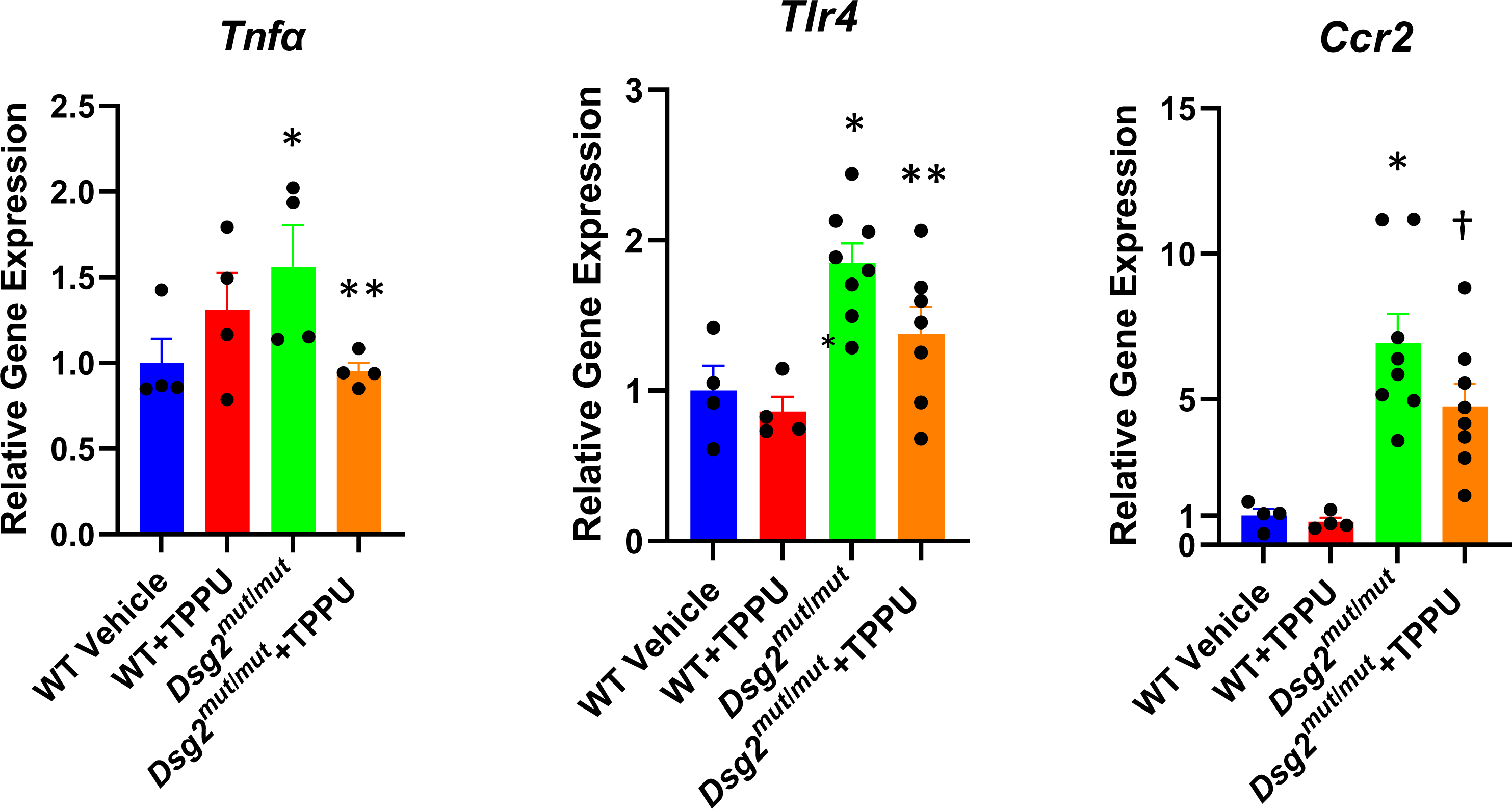
Effects of 1-trifluoro-methoxy-phenyl-3-(1-propionylpiperidin-4-yl) urea (TPPU) on expression of *Tnfα*, *Tlr4* and *Ccr2* in hearts of wildtype (WT) and *Dsg2^mut/mut^* mice treated with TPPU or vehicle. Gene expression values in wildtype samples treated with vehicle were normalized to 1; values in all other groups are shown as relative levels; * p<0.05 vs. WT Veh; ** p<0.05 vs. *Dsg2^mut/mut^* mice; † p=0.0579 vs. *Dsg2^mut/mut^* mice, all by multiple comparisons ANOVA.

## DISCUSSION

Inflammation is usually a self-limited process, designed to eliminate the cause of injury and restore homeostasis.^8,21^ If left uncontrolled, however, inflammation can cause progressive tissue damage. Indeed, diverse diseases such as sepsis, acute respiratory distress syndrome,cancer and COVID-19 are driven by unresolved inflammation.^22,23^ Resolution of inflammation is an active regulated process, orchestrated in part by various endogenous specialized pro-resolving mediators (SPMs) synthesized from ω-3 polyunsaturated fatty acids (PUFAs) by lipoxygenase, cyclooxygenase and/or cytochrome P450 mono-oxygenase enzymes (CYP450s).^24^ EpFAs not only increase SPM production but are also direct pro-resolving mediators.^25,26^ SPMs also promote resolution of pathological endoplasmic reticulum (ER)-stress, a hallmark of unresolved inflammation.^9^ However, many pro-resolving EpFAs are short-lived. For example, EETs are rapidly metabolized by sEH to their corresponding diols, including dihydroxyeicosatrienoic (DiHETES) and dihydroxyoctadecenoic (DiHOMES) acids, which may exert pro-inflammatory effects.^11,12,25^ Small molecule inhibitors of sEH stabilize levels of pro-resolving EpFAs and reduce tissue injury in various animal models.^25–28^ sEH inhibitors shift arachidonic acid metabolic cascades from a pattern of initiation of inflammation to one of resolution.^29^ Importantly, both SPMs and EpFAs, synergized by inhibition of sEH, potently down-regulate NFκB pathways,^29,30^ which are persistently activated in cardiac myocytes in a cell-autonomous fashion in ACM.^3–7^

Increasing evidence suggests that chronic inflammation in ACM is the result of deficient resolution. The consistent presence of inflammatory infiltrates in the hearts of ACM patients obviously implicates immune mechanisms.^1,2^ Moreover, ACM patients have elevated circulating levels of cytokines and their cardiac myocytes express multiple pro-inflammatory mediators.^31^ Recurrent bouts of inflammation (rigorously defined as troponin elevation with normal coronary arteries, and typical ^18^F-fluoro-deoxyglucose PET findings) occur in ACM patients with variants in *DSP*, the gene for the desmosomal protein desmoplakin.^32^

Here, we provide new evidence that 1) ACM is driven by a persistent innate immune response associated with reduced levels of pro-resolving bioactive lipid mediators; 2) the pro-resolving EpFA 14,15-EET can reverse disease features in an *in vitro* model of ACM; and 3) inhibition of the sEH enzyme mitigates the disease phenotype in a well characterized animal model of ACM. sEH inhibitors are currently being evaluated in phase 1 clinical trials. sEH inhibitors are in clinical development for hypertension,^33^ chronic obstructive pulmonary disease,^34^ chronic pain and other conditions.^14,35^ Dual COX-2/sEH inhibitors are also currently in clinical development for multiple inflammatory diseases.^36,37^ Should these small molecule drug candidates be found to be safe and free of toxic side-effects, our observations suggest that they may be of benefit in patients with ACM.

## Acknowledgments

This work was supported by NIH grant R01HL148348 (JES) and 1R01CA276107-01A1 DP, and grants from Credit Unions Kids at Heart and the Carter Joseph Buckley Pediatric Brain Tumor Fund (DP). Partial support was provided by NIH – NIEHS (RIVER Award) R35 ES030443- 01, NIH-NINDS U54 NS127758 (Counter Act Program), and NIH – NIEHS (Superfund Award) P42 ES004699 (all to BDH). Additional support came from a Washington University in St. Louis Rheumatic Diseases Research Resource-Based Center grant (NIH P30AR073752, KL), a National Institutes of Health grant (R35 HL161185, KL), a Leducq Foundation Network grant (#20CVD02, KL, a Burroughs Welcome Fund grant (1014782, KL), a Children’s Discovery Institute of Washington University and St. Louis Children’s Hospital grant (CH-II-2015-462, CH-II-2017-628, PM-LI-2019-829, KL), a Foundation of Barnes-Jewish Hospital grant (8038- 88, KL), and gifts from Washington University School of Medicine (KL). V. Penna was supported by a National Institutesof Health grant (5T32AI007163-44). Additional support came from the British Heart Foundation (PG/18/27/33616; CBB, AA).

